# The difficulty in numerical computation impacts motor decisions in a Stop Signal task

**DOI:** 10.1101/2025.07.01.662562

**Authors:** Isabel Beatrice Marc, Valentina Giuffrida, Mariella Segreti, Ann Paul, Sabrina Fagioli, Pierpaolo Pani, Stefano Ferraina, Emiliano Brunamonti

## Abstract

The proper interpretation of environmental information is necessary for effective decision-making. The resulting cognitive burden may affect the entire process if interpretation is not instantaneous. In this study, we investigated how numerical distance (ND), a measure of cognitive demand in numerical comparisons, influences movement initiation and inhibition. To this end, 32 participants completed a novel numerical comparison Stop-Signal Task (NC-SST), in which the cognitive demand of each trial was manipulated by varying the ND between pairs of numbers in both Go and Stop signals. Participants were required to initiate or stop a movement if an upcoming number was higher or smaller than the one presented previously. Results showed that larger NDs (i.e., easier comparisons) facilitated faster and more accurate responses during movement initiation and enhanced stopping performance. Using a generalized drift-diffusion model, we found that drift rates increased with ND and were modulated by the spatial location of numerical stimuli, consistent with a left-to-right space number association. A generalized linear mixed-effects model further revealed that Go process parameters, particularly the drift rate, strongly predicted successful stopping and interacted with Stop ND and Stop signal delay (SSD). These findings demonstrate that higher cognitive load impairs both movement initiation and inhibition, and that motor decisions result from the integration of cognitive information onto perceptual features, extending the classical race model framework.

## INTRODUCTION

We frequently have to decide whether to stop an action responding to something unexpected or to adapt to changing environmental demands. An experimental paradigm that probes this behavior is the Stop-Signal task (SST; Logan & Cowan, 1984; Aron & Verbruggen, 2008; Bissett & Logan, 2011). In the SST, participants usually have to generate a motor response as quickly as possible to a “Go signal” or cancel the planned movement after the presentation of a “Stop signal”. In the context of the SST, the theoretical framework of the ‘Horse-Race Model’describes a competition (race) between response inhibition elicited by the Stop signal and response initiation triggered by the Go signal. Within this framework, the winner of the race determines the behavioral outcome, such as whether the response continues or stops. It has been shown that various factors, such as motivation (Leotti & Wager, 2010; Boehler et al., 2012, 2014; Giamundo et al., 2021; Giuffrida et al., 2023; Verbruggen & McLaren, 2018), attention (Verbruggen et al., 2014; Hilt & Cardilicchio, 2020; Haque et al., 2024), and the perceptual characteristics of the stimuli (Montanari et al., 2017; Middlebrooks et al., 2017, 2019; Pani et al., 2018; Marc et al., 2023), can influence performance when cancelling the planned movement is required.

The influence of cognitive load on motor control during visuomotor transformation has been less thoroughly investigated. Some experimental contexts have explored this decision difficulty using tasks that require comparisons between rank-ordered items or numerical pairs (Merritt & Terrace, 2011; Brunamonti et al., 2012; Brunamonti et al., 2016; Jensen et al., 2017; Mione et al., 2020). These experiments have demonstrated that the difference between the ranks or between the numerical values of the stimuli influences decision-making accuracy and latency (see Ramawat et al., 2023 for a review).

The spatial-numerical associations construct accounts for difficulty in numerical representation and comparisons. This abstract representation is spatially oriented from left to right with single positions partially overlapping (Dehaene et al., 1993; Dehaene, 1997; Hubbard et al., 2005; Umiltà et al., 2009, see Fischer & Shaki, 2014 for a review). According to studies demonstrating an association between numerical and spatial information, one determines whether 7 is greater than 5 by comparing their positions on this internal mental representation, where small numbers (e.g., 1) are associated with the left side of the space and larger numbers (e.g., 9) with the right (Longo & Lourenco, 2007). A phenomenon observed during numerical comparisons is the numerical distance (ND) effect, which modulates accuracy and reaction time depending on the distance in the internal representation (Moyer & Landauer, 1967; Hubbard et al., 2008). When two numbers are relatively far apart, participants respond faster and accurately than when they are closer (Izard & Dehaene, 2008; Holloway & Ansari, 2009).

The study investigates the effect of the ND on motor control, with the underlying hypothesis that assigning a cognitive load onto perceptual detection modulates motor decisions. Here we developed a numerical comparison Stop Signal task (NC-SST), where participants evaluated numerical pairs, which integrates ND into both Go and Stop signals. The behavioural results demonstrated that ND, manipulated across 5 levels, had a significant effect not only on movement initiation but also on inhibition, reflecting the interplay between cognitive load and motor control. With this design, we could also test whether differences in cognitive load would be reflected in the underlying dynamics of decision-making processes. We used the drift-diffusion model (DDM; Ratcliff, 1978; Shinn et al., 2020) for this purpose, which views decision-making as the process of accumulation of information over time. Based on this framework, we hypothesised that larger ND, reflecting easier comparisons, would be associated with faster drift rates, leading to quicker response initiation. To further examine how the accumulation of Go-related evidence influences movement inhibition, we applied a generalized linear mixed-effects model (GLMM) to predict stopping success on a trial-by-trial basis. This analysis tested whether faster Go Drifts impaired the ability to stop, and whether this effect was modulated by the characteristics of the Stop signal, such as its ND from the Go signal, as supported by our results.

## 2. METHODS

### 2.1 Participants

Thirty-two healthy participants (27 females and 5 males, mean age 25 ± 4,4) were recruited for the study. According to the Edinburgh Handedness Inventory (Oldfield, 1971), all participants were tested for manual dominance, and there were 1 left-handed and 31 right-handed participants. Participants had normal or corrected vision without any known neurological or psychiatric conditions. All procedures followed the Declaration of Helsinki and after obtaining written informed consent from each participant. The procedure was approved by the Ethics Committee of “Roma Tre” University.

### 2.2 Experimental Procedure and the NC-SST

Participants were seated in a darkened, sound-attenuated room facing a monitor (15.6 inches) on a laptop computer that was comfortably situated 50-60 centimetres away. Under different experimental conditions, participants performed a modified version of the SST, in which the Go and the Stop signals were represented by the outcome of a mental computation, rather than the encoding of a perceptual stimulus. Each participant held a mouse while resting their arm on a table. Each trial started when both effectors (the index and middle fingers) simultaneously pressed both mouse buttons to reach their initial positions.

Every time the buttons were pressed, a Wait Signal would show two identical numbers, such as 5-5, 283 pixels to the left and right of the center of the screen. Each of the numbers was 20 × 28 pixels in size and white, and the overall resolution of each image was as high as the screen resolution with a black background (Figure 1**a-c**). In Go trials (50% of the trials), one of the two numbers increased (for example, 5-7 from 5-5) following a variable waiting time (500–1000 ms; in steps of 50 ms).

**Figure 1.**
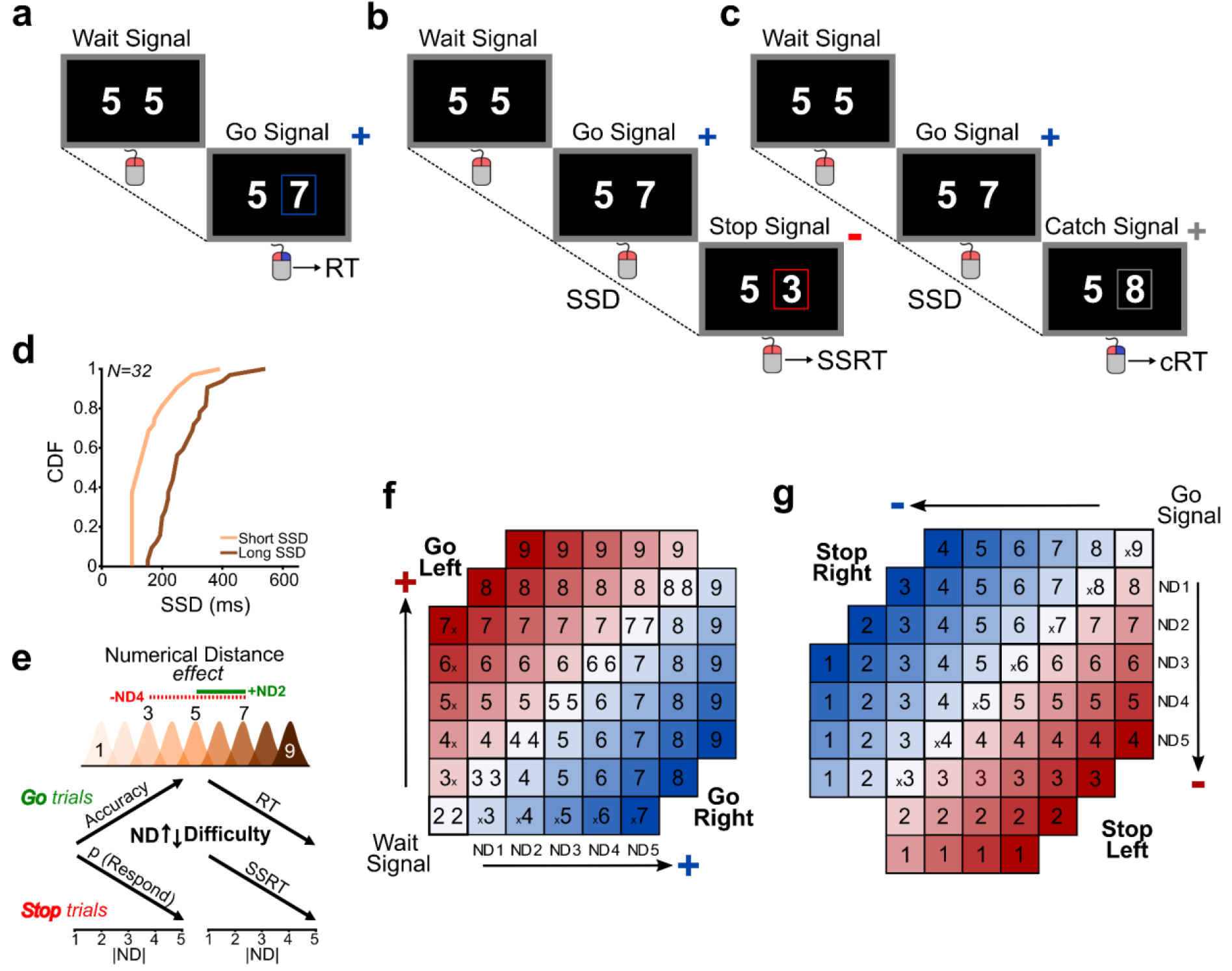
Illustration of the experimental task and the manipulation of the degree of difficulty in the different conditions. **a-c** *Time course of the task’s three different trial types*. **a** In Go trials, the mouse button that needed to be released was indicated by an increasing value of the number on one side of the screen (Go Signal). **b** In Stop trials, a reduction in value at the same screen position following the Go Signal indicates to stop the engaged movement. **c** In Catch trials, a second increase in value indicated to continue and complete the movement. **d** The cumulative distribution function (CDF) displaying participants’SSD distributions (*N*=32; short SSD= light orange; long SSD= brown). **e** The hypothesized spatial representation of the numbers hierarchy. The ND effect is illustrated through distributions, where lower NDs cause greater overlap in mental representation, which makes comparisons harder, while higher NDs reduce overlap, which makes comparisons easier. Hypothesized behavioral outputs in Go and Stop trials. In Go trials, Accuracy is increasing while RTs are decreasing, while in Stop trials, p(Respond) to the Stop Signal and SSRT are decreasing with greater ND. **f-g** *Schematic representation of all combinations in Go and Stop trials*. **f** All Wait Signals are presented in the diagonal squares, same number on both sides of the screen. The following Go Signal was represented by an increase in the value of one of the two numbers, either on the Left (red color scales) or on the Right (blue color scales) as a function of NDs. **g** In Stop trials, diagonal squares represent the Go Signals, and the following Stop Signal was represented by a decrease in the value as a function of ND from the Go signal. RT, Non-Stop Reaction Time; SSD, Stop Signal Delay; SSRT, Stop Signal Reaction Time; cRT, Catch Reaction Time; ND, Numerical Distance.

This change served as the Go Signal, which instructed participants to release the mouse button congruent with the spatial location of the number that changed value on the monitor (Figure 1**a**; Figure 1**f** shows all combinations of Wait and Go Signals). In Stop trials (25% of the trials), after a variable amount of time (SSD; Stop Signal Delay) from the Go Signal, a Stop Signal with a smaller value, such as 5-3 following 5-7 (Figure 1**b**; Figure 1**g** shows all combinations of Go and Stop Signals), placed the Go Signal. In this case, the Stop Signal instructed participants to inhibit the movement triggered by the previous pairs of numbers. To ensure that participants focused on the numerical comparison to be made rather than simply reacting to any changes after the Go signal, we included Catch trials (remaining trials) alongside Stop trials (Figure 1**c**). In these trials, the stimulus occurring after the Go signal was based on a change to a higher number, such as 5-8, instructing the participants to continue their movement. These trials served as a control condition and were not included in the behavioral or modeling analyses. First, participants experienced the three trial types (Go, Stop, and Catch) in a familiarisation block (~ 100 trials).

During this familiarization block, a tracking procedure dynamically adjusted the duration of the SSD in steps of 50 ms based on the accuracy of the previous Stop trial: initial value was set to 200 ms increasing by one step (+50 ms) if the trial was executed correctly and decreasing by one step (−50 ms) if executed incorrectly. Catch trials were presented using the most recently computed SSD value, ensuring consistency with the last adjustment made during the Stop trial. We then computed the average SSD for each participant during the familiarization block and defined two difficulty conditions based on this value, in line with the horse-race model, a *short SSD* condition (average SSD − 100 ms; population mean: 161 ± 71 ms) and a *long SSD* condition (average SSD + 50 ms; population mean: 271 ± 86 ms). Figure 1**e** displays the distribution of these values among participants, which were utilised in the test phase that followed. In addition to the difficulties introduced by the length of the SSDs when interpreting the visual stimuli, as either a Stop or a Catch signal, we varied the ND in the paired number comparison to further manipulate the complexity of cognitive operations needed in the response, from small distances to larger distances (e.g., 4-5: ND1 or 4-9: ND5; for an overview of combinations see Figure 1 **f-g**). Furthermore, to avoid potential bias brought on by participant expectations, NDs were maintained constant (using only ND1 or ND2) during the testing phase of Catch trials. For instance, because of the combined effect of the increased ND, a high-value number on the Go signal would be less likely to indicate a Catch trial and more likely to signal a Stop trial. During this phase, each participant performed the task divided into 4 blocks with a 10-minute pause between each block (~ 1500 trials). Two separate auditory feedbacks were used to distinguish between trials that were completed successfully and those that were not. The presentation of stimuli and collection of behavioral events were handled with MATLAB® R2021b (www.mathworks.com), using the Psychophysics Toolbox Version 3 feature set (www.psychtoolbox.org).

### 2.3 Variables Estimation and Data Analysis

The current study aimed to assess how the difficulty of interpreting NDs among numbers influenced both movement initiation and inhibition in response to perceptual signals. To achieve this, we examined how participants’performance was influenced by NDs (from 1 to 5) across both Go and Stop trials. In Go trials, we also assessed the influence of the Spatial Location (SL; Left/Right) of the higher number, while in Stop trials, we evaluated the effect of inhibition timing by manipulating SSDs (Short/Long). Using a 2×5 within-subjects design, we conducted two-way repeated measures ANOVAs to examine: (1) Accuracy, which represents the proportion of correct responses to the Go Signal, providing an indicator of overall performance; (2) Go RTs, measured as the time between the presentation of the Go signal and the button release. Omission errors (lack of movement initiation) and commission errors (initiating movement in the wrong SL indicated by the Go Signal were excluded; and (3) Go RTs Variability, measured as the standard deviation of the Go RT distribution, assessed the consistency of participants’RTs across trials. In addition to these measures of movement initiation, to assess movement inhibition, we also examined: (1) p(Respond) to the Stop signal, the probability of responding on Stop Trials, which reflected participants’ability to inhibit their response when a Stop Signal was presented; and (2) Stop Signal RT (SSRT), which is a nonparametric estimate of the time participants required to successfully inhibit the response after the Stop signal, computed through the integrative method (Verbruggen et al., 2019). For each ND, this was computed using trials from both SSDs to get an accurate estimate (50> Stop trials). Each participant’s performance was checked for compliance with the independence assumption of the ‘horse-race model’, as indicated in (Verbruggen et al., 2019) for the reliable estimate of the SSRT. We considered the independence assumption respected if the average Stop Error RTs was lower than the average Go RTs (De Jong et al., 1995; Verbruggen & Logan, 2009a). These averages were compared using paired ***t****-*tests; the relevant variables were computed for each participant separately and for each ND. Custom MATLAB® R2022b (www.mathworks.com) functions were implemented for data processing and analysis.

### 2.4 Fitting the generalized drift-diffusion and generalized linear mixed-effects models

***GDDM***. Using a generalized drift-diffusion models toolbox (PyDDM; Shinn et al., 2020), we applied the generalized drift-diffusion model to participants’Go RTs distributions to determine the computational strategies employed across ND. The model was originally developed to describe perceptual decision-making (Ratcliff & McKoon, 2008), but it has since been expanded to include symbolic manipulation and numerical cognition (Dehaene, 2009; Ratcliff et al., 2015; Park & Starns, 2015; Krajcsi et al., 2016; Dix & Li, 2020). The GDDM conceptualizes decision-making as a noisy, continuous process in which information accumulates over time, where a choice is made when the decision variable *(dx)* reaches one of two decision boundaries *“B”*, which in our case correspond to accuracy (Correct vs. Error) or target choices (Left vs. Right), based on model hypothesis tested. To flexibly capture these dynamics, we implemented four models of increasing complexity, each building upon a basic drift diffusion framework - the null model. For all the tested models, time resolution (*dt*) was set to 0.01 seconds, and the overall duration was set to 1.5s, corresponding to the upper RT limit to respond in Go trials. The Null model captures only the basic decision dynamic, not including any influence of the stimulus characteristic or decision context. The evolution of the *dx* is governed by: *dx* = *ν*∗*dt*+*dW*

It assumes a constant drift parameter *“v”* (=10), symmetric decision boundaries (= 3) set as “Correct” for upper bound and “Error” for lower bound, a fitted non-decision time (= 0.1s), and *“W”* is momentary noise (std=1).

*Model 1 (“Linear”)*. Building upon the Null model, we added a condition-dependent *“v”* to test whether it varies linearly with ND, where the evolution of the*“dx”* is governed by: *dx*=[*ν*∗*ND*]∗*dt*+*dW*

All parameters are estimated to vary freely within a range (*ν* = −10 to +10, *B* = 0.1 to 3; and non-decision time = 0.05s to 0.4s).

*Model 2 (“LinearwithLeak”)* improves Model 1 by adding the possibility that imperfect integration “*leak*” of evidence can happen, modeling memory decay where *dx* is governed by:

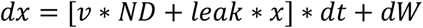

The *“leak”* parameter (−10 to 10) is a constant that acts as a multiplier modulating how accumulated evidence*“dx”* changes over time. A negative leak causes the evidence to decay towards zero, while a positive leak makes it grow exponentially.

*Model 3 (“LinearwithSLbias”)* offers an extension to Model 1 by introducing a spatial location bias (SL bias) toward one of the two target choices (Left = upper bound vs. Right = lower bound), spatial biases not accounted for by the ND alone, where*“dx”* is governed by:

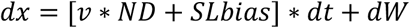

The SL bias is a free parameter (−5 to +5) capturing a consistent bias toward the Left or Right *“B”. Model 4 (“LinearwithSLbiasLeak”)* integrates upon Models 2 and 3 to offer a more complete description of the decision process. It assumes that the “*v”* varies linearly with ND, includes SL bias, and adds a *leak* term to capture imperfect evidence integration. The *“dx”* evolution evolves according to:

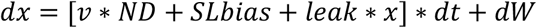

This model provides a flexible framework to test how the ND effect, SL bias, and evidence leak jointly contribute to decision behavior.

***GLMM***. To explore how the processing of Go and Stop signals interacted at the single-trial level, in a subgroup of 17 participants, in which Go signal ND information was available, we encoded for each trial the ND eliciting the Stop process, depending on the one that in the same trial triggered the Go process. We then fitted a generalized linear mixed-effects logistic regression to examine how trial-wise parameters predict successful inhibition. The binary outcome variable S_ij_∈{0,1}, indicating successful (1) or failed (0) inhibition for each trial *i* and participant *j*, was modeled as a function of Go Drift, Stop SSD (Short/Long), Stop ND, and Stop SLbias (Left/Right), along with relevant interactions. The mixed-effects structure included random intercepts and random slopes for Go Drift by participant, to account for individual variability in baseline stopping performance and sensitivity to Go evidence. The model specification: *Sij*=*β*0+*β*1*GoDrift*+*β*2*StopSSD*+*β*3*StopND*+*β*4*StopSLbias*+*β*5(*GoDrift*StopSSD*)+*β*6(*GoDrift*StopND*)+*β*7(*StopND*StopSSD*) +*u*0*j*+*u*1*j GoDrift* where β0 is the intercept, β1 to β7 are fixed effect coefficients, and u_0j_, u_1j_ are participant-specific random intercepts and slopes.

## 3. RESULTS

### 3.1 Movement initiation is influenced by ND

We assessed the influence of spatial location (SL; Left/Right) and ND (1-5) during movement initiation on Accuracy (Figure 2**a**) and found a significant increase with increasing ND (two-way repeated measures ANOVA; F(_4,124_)=7.19, p < .001) but no significant main effect of the SL (F(_1,31_)=0.60, p= .13) or significant interaction (F(_4,124_)=2.81, p= .71). We then examined whether Go RTs and their Variability were influenced by SL and ND (Figure 2**b-c**).

**Figure 2.**
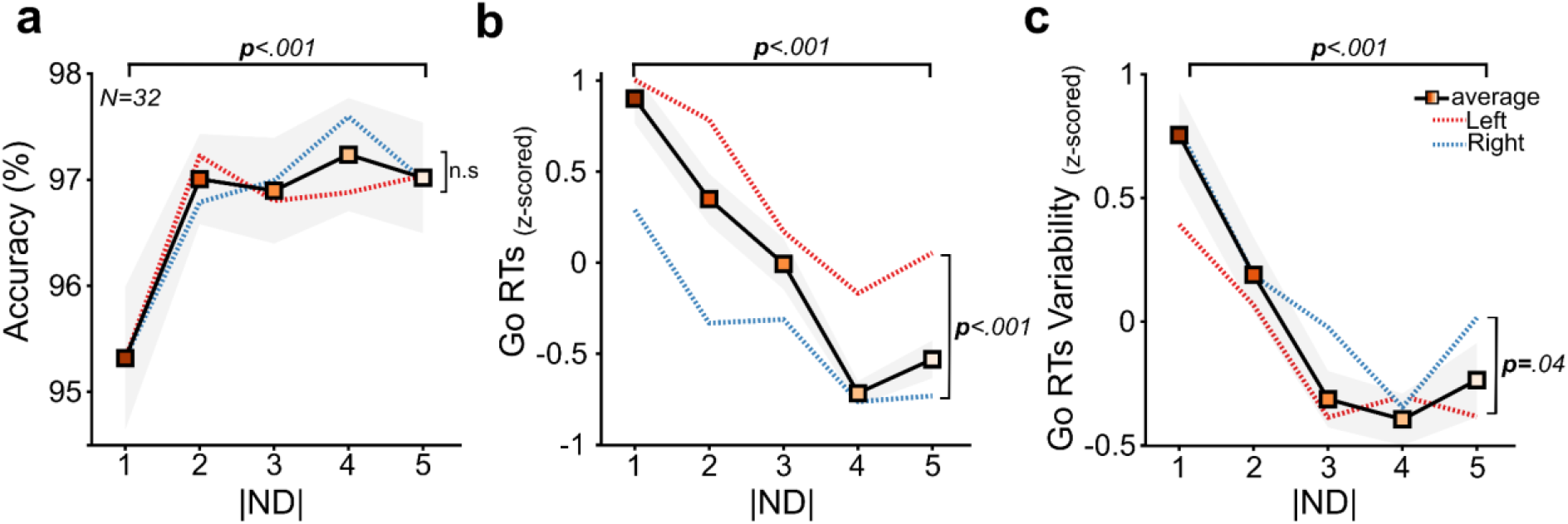
Performance in Go trials. **a)** Accuracy, **b)** Go RTs (z-scored), and **c)** Go RTs Variability (z-scored). The analysis considered the representation of both the Left (depicted by a dashed red line) and Right (indicated by a dashed blue line) SL of the Go Signal presentation. Shaded areas and lines illustrated the averaged values between left and right for each ND. Grey shaded areas are ±1 SEM. average = averaged across SL

Go RTs were significantly faster if the higher-number was shown on the Right (mean: Left= + 0.36; Right = −0.36) (two-way repeated measures ANOVA; F(_1,31_)=19.50, p < .001) and as comparison became easier (F(_4,124_)=25.76, p < .001), with no significant interaction (F(_4,124_)=2.05, p =.05). Additionally, RTs Variability was higher when the higher-number was displayed on the Right (mean: Right= + 0.12; Left = −0.12) (two-way repeated measures ANOVA; F(_1,31_)=4.26, p = .04), but decreased significantly as NDs increased (F(_4,124_)=11.87, p < .001), with no significant interaction was found (F(_4,124_)=0.97, p = .42).

Our findings suggest that ND influences movement initiations during Go trials, resulting in increased Accuracy with faster and more consistent RTs when executing movements signaled by the higher number. Additionally, RTs were faster but more variable when a higher number was displayed on the Right side.

### 3.2 The difficulty in properly inhibiting a movement was influenced by NDs and SSDs

To evaluate the effect of ND during Stop trials on the ability to inhibit a movement, we first needed to assess whether participants respected the race model’s independence assumptions, as violations would preclude a reliable estimation of the SSRT. We used paired ***t***-tests to assess context independence at the group level by comparing Stop Error RTs with Go RTs for each ND (Table 1).

**Table 1.** Assessment of context independence across ND. Average Stop Error RTs and Go RTs for each ND, along with standard deviation and results of paired ***t***-tests comparing the two variables at a population level.

| ND | Stop Error RT<sub>(ms)</sub> | Go RT<sub>(ms)</sub> | *t-value*<sub>(31)</sub> | *p-value* |
| --- | --- | --- | --- | --- |
| 1 | 550 (±135) | 571 (±87) | -0.80 | 0.43 |
| 2 | 531 (±127) | 557 (±79) | -0.97 | 0.33 |
| 3 | 498 (±131) | 551 (±80) | -1.93 | 0.06 |
| 4 | 490 (±148) | 543 (±77) | -1.92 | 0.06 |
| 5 | 483 (±141) | 545 (±77) | -2.50 | 0.01 |

In line with the predictions of the race model, the overall trend indicated that Stop Error RTs were shorter than Go RTs, even though a significant difference was observed only at ND5, indicating that context independence held more consistently for easier comparisons. This pattern suggested that a higher number of violations might be linked to more challenging comparisons. We next assessed context independence at the individual level and observed that as ND increased, the number of participants violating the model’s assumption decreased (Figure 3**a**). We quantified these violations and found that they occurred most frequently at smaller NDs (Figure 3**b**; small insert panel: ND1= 18; ND2= 15; ND3= 12; ND4= 12; and ND5 = 8). We next examined the p(Respond) to the Stop signal, quantifying the proportion of trials in which participants failed to inhibit their response, as this measure offers a general indicator of stopping performance and is independent of the model’s assumptions (Figure 3**c**). Results revealed significant main effects of SSD (two-way repeated ANOVA; F(_1,31_) = 46.42, p < .001) and ND (F(_4,124_) = 23.73, p < .001), indicating that stopping was less successful at longer SSDs and lower ND. No significant interaction was observed (F(_4,124_) = 2.27, p = .06).

**Figure 3.**
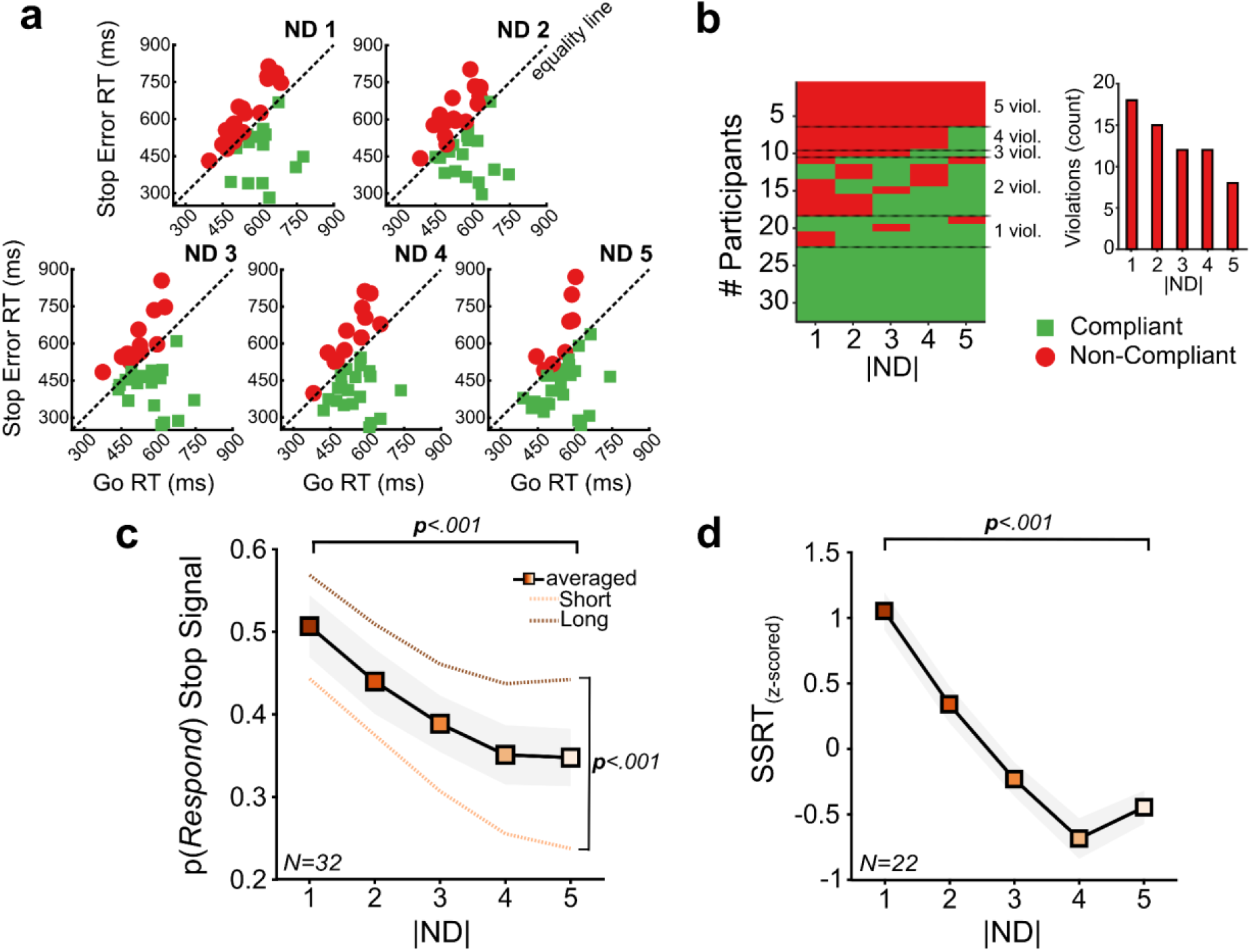
Assessment of movement inhibition. **a** Horse-Race model’s assumptions evaluation for each ND. Each participant is represented by a colored marker; green squares indicate a Compliant participant with model assumptions, while red dots indicate Non-Compliant participants. The equality line is displayed. **b** Quantification of participants violating the model’s assumptions across NDs. The small insert panel counts the number of participants violating the model’s assumptions. **c-d** Influence of ND on p(Respond) to the Stop Signal and SSRTs (z-scored) depicted for both Short (depicted by a dashed light orange line) and Long (depicted by a dashed brown line) SSDs. Grey shaded areas are ±1 SEM; average = averaged across SSD.

Finally, we estimated SSRT for a subset of participants who largely complied with the model’s assumptions. We applied a threshold to define this subgroup, and participants were excluded if they violated the independence assumption in more than 3ND. This resulted in a subgroup of Non-Compliant participants (*N* = 10) and the estimation of SSRT for a Compliant group (*N* = 22) that did not exhibit a consistent pattern of violations across ND (Figure 3**d**). The analysis revealed significant main effects of ND (one-way repeated ANOVA *F*(_4,84_)= 18.17, *p* < .001). These findings indicate that the difficulty of comparisons posed by the effects ND affects the validity of the assumptions of the race model and influences stopping performance.

### 3.3 Go Drift rates underlie Go RT differences and predict Successful Stopping performance

To assess GDDM model performance, we used the Bayesian Information Criterion (BIC). Among the candidate models, we found that Model 4 provided the best-fitting model to the data (Figure 4**a**), indicating that decision-making performance was best explained by variations in Go drift rates across ND, SLbias, and the leak term capturing imperfect evidence integration. In addition to yielding the lowest BIC, Model 4 also had the highest (least negative) log-likelihood, confirming that its superior fit was not simply due to a higher number of parameters. Average parameter estimates for each model are reported in Table 2.

**Table 2.** Average fitted parameters across participants for each model by the GDDM. Model 1. *“Linear”*; Model 2 *“LinearwithLeak”*; Model 3 *“LinearwithSLbias”*; Model 4 “*LinearwithSLbiasLeak”*.

| Model | Drift | SLbias | Leak | Non-decision (ms) | BIC |
| --- | --- | --- | --- | --- | --- |
| 1 | .80 (± .11) | - | - | 317 (± 79) | - 415.7 (± 567.8) |
| 2 | -.22 (± .31) | - | 9.05 (± 1.39) | 231 (± 85) | + 32.4 (± 408.1) |
| 3 | .84 (± .10) | - .08 (± .13) | - | 314 (± 79) | - 599.7 (± 442.1) |
| 4 | .91 (± .14) | - .15 (± .22) | 6.34 (± 1.82) | 232 (± 81) | - 766.4 (± 448.8) |

**Figure 4.**
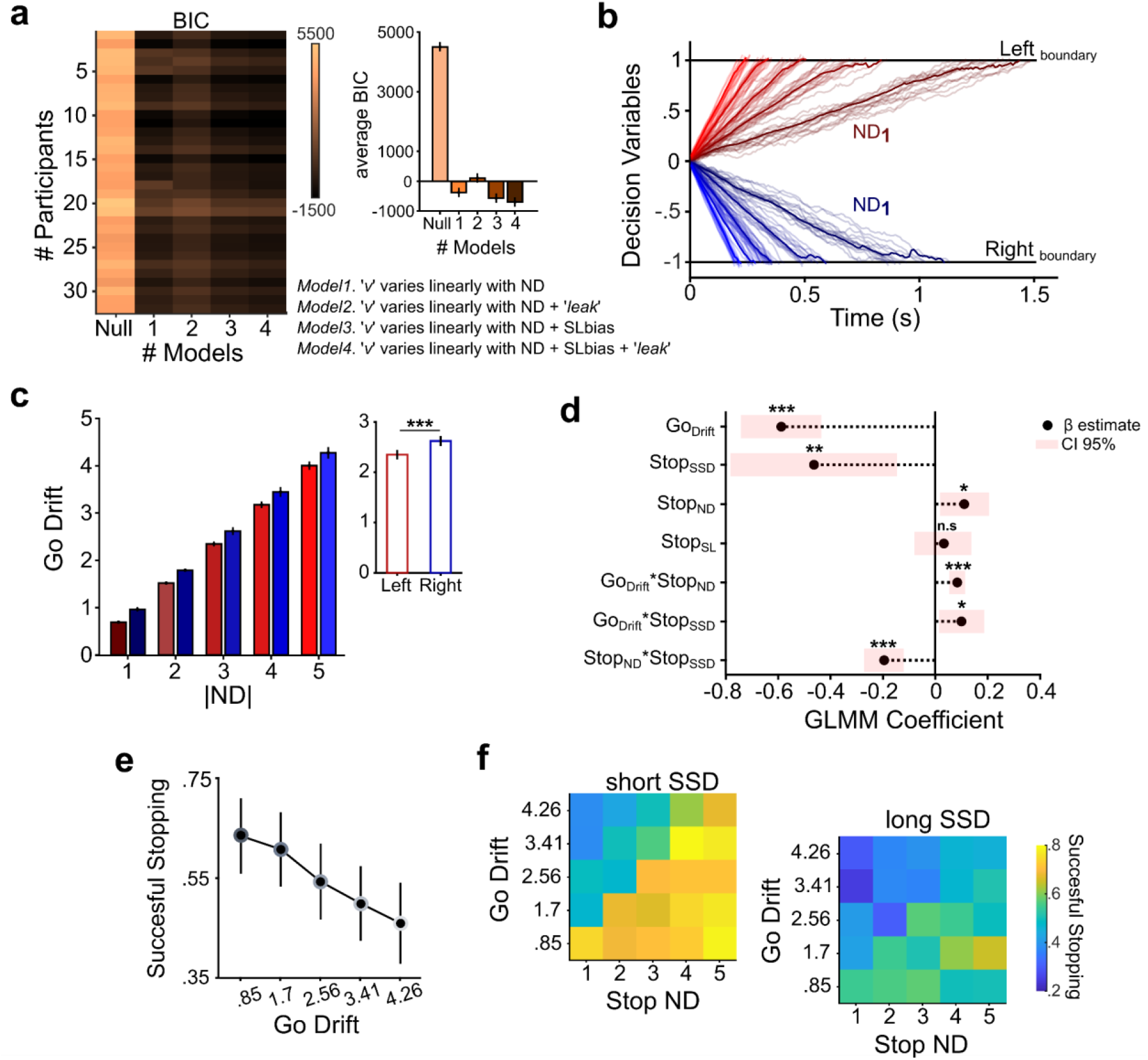
Effects explained with the GDDM and GLMM. **A** BIC values for each fitted GDDM model (from Null to Model 4), with each row representing a participant and an insert panel showing group averages ±1 SEM. Model 4 shows the best fit overall. **b** Simulated trajectories from the GDDM for a representative participant, using the Go Drift rates from the best-fitting model. Each colored trace represents a single simulated trial under a given ND and SL (Left = red shades, Right = blue shades). Darker solid lines show the average across trials; black horizontal lines mark decision boundaries. **c** Bar plot showing average Go drifts by ND and SL, with error bars indicating ±1 SEM. The small insert panel shows the significant overall difference between Left and Right SL, with generally higher Go drift rates for the Right SL, suggesting lateralization in processing speed. **d** GLMM Coefficient β estimate with a 95% confidence interval. **e** Average successful stopping probability decreases as the Go Drift rate increases; higher drift rates triggered by the Go signal with larger ND values correspond to lower successful stopping probability. Error bars represent ±1 SEM. **f** Heatmaps showing successful stopping probability as a function of Go Drift (y-axis) and Stop ND (x-axis), separately for short SSD (**f**) and long SSD (**g**). The effect of Go Drift on successful stopping is mitigated by Stop ND: higher Stop ND distances lead to increased stopping probability, whereas lower Stop ND distances reduce stopping probability. Additionally, shorter SSDs (left panel) show generally higher stopping probabilities than longer SSDs (right panel), illustrating the impact of SSD on inhibition success. Colorbar indicates the successful stopping probability. Asterisks indicate significance (^*\**^< .05, ^**\*\***^< .01, ^***\*\*\****^ < .001).

To visualize these dynamics, Figure 4**b** shows simulated decision trajectories generated using Go Drift rates from the best-fitting model for a representative participant, highlighting the influence of both ND and SL bias.

The trajectories show faster evidence accumulation toward the lower Right boundary, reflecting the rightward SL bias (−0.13 ± .20), while accumulation toward the upper Left boundary is comparatively slower. Analysis of the fitted Go Drift rates revealed an increase with ND (ND1= 0.82 ±.11; ND2= 1.65 ±.22; ND3= 2.48 ±.34; ND4= 3.31 ±.45; ND5= 4.13 ±.57), showing faster evidence accumulation for larger NDs. Furthermore, there were significantly higher Go Drifts towards the Right SL (small insert panel; paired ***t***-test: ***t***_**(159)**_= −8.48, *p*<.001) (Figure 4**c**).

Building on these findings, we next examined whether Go Drift rates influenced trial-wise stopping performance. Using a GLMM, we predicted the probability of successful stopping based on parameters derived from Stop trials (see Methods for details and Figure 4**d**). The analysis revealed that higher Go Drift values were significantly associated with lower stopping success (β_1_= –0.59, *p* < .001), supporting the hypothesis that stronger Go-related evidence accumulation impairs inhibition. This relationship was also observed in the empirical data, where higher Go Drift corresponded with lower stopping success (Figure 4**e**). As expected, longer SSDs were linked to decreased stopping success (β_2_= –0.46, *p* = .004), while greater Stop ND (between Go and Stop signals) predicted improved stopping performance (β_3_= +0.11, *p* = .02). This suggests that increased ND distinction between Go and Stop signals facilitates inhibitory control. In contrast, Stop SL showed no significant effect after controlling for other predictors (β4= +0.03, *p* = .60). We also observed a significant interaction between Go Drift and SSD (β_5_ = +0.10, p = .02), indicating that Stop trials with both high Go Drift and long SSDs were associated with particularly poor stopping performance. However, this detrimental effect of Go Drift was mitigated when Stop ND was high, as shown by a significant Go Drift × Stop ND interaction (β_6_= +0.08, *p* < .001). A further interaction between Stop ND and SSD (β_7_ = –0.20, *p* < .001) revealed that the benefits of high Stop ND were especially pronounced at shorter SSDs (Figure 4**f**). Overall, these results indicate that stopping success depends not only on the individual properties of Go and Stop processes but also on their interactions. Specifically, inhibitory control is shaped by the strength of Go evidence (Go Drift), the temporal window for inhibition (SSD), and the distinctiveness of the Stop signal (ND), all of which jointly determine stopping efficiency on a trial-by-trial basis.

## DISCUSSION

Our study shows that both movement initiation and inhibition were influenced by the cognitive load and the difficulty of interpreting the information provided by the stimuli. Different levels of cognitive load were introduced by variations in the ND, even though the visual appearance of the stimuli remained perceptually unambiguous across trials. Our results suggest that motor decisions are influenced by cognitive processing, not only by stimulus-driven visuomotor transformations, offering new insights into how decision-making integrates both cognitive and perceptual content. Previous studies focused primarily on how perceptual difficulty influences motor performance during Go trials. Pressing buttons or saccadic responses have been prompted by the spatial location of visual stimuli (Logan, 1981; Logan & Irwin, 2000). While other studies varied the visual similarity of alphabetic letters (Osman et al., 1986), with varying degrees of distinguishability, such as “G vs. X” or “I vs. i” for relatively easy or difficult comparisons, showing that increased ambiguity increases the latency of the responses. Middlebrooks and Schall (2013) similarly varied the proportion of colours in a checkerboard search array that cued leftward or rightward saccades, so altering target discriminability. They found that saccade latency decreased as color dominance increased. These results prove that faster movements are facilitated by the perceptual process. On the other hand, a different line of research concentrated on how the salience of the Stop signal affects movement inhibition, showing that reducing the perceptual discriminability results in slower inhibition (Van Der Schoot et al., 2005; Morein-Zamir & Kingstone, 2006; Montanari et al., 2017; Middlebrooks et al., 2019). In Montanari et al. (2017), participants initiated reaching movements based on a colored Go signal and stopped them depending on a color change in the Stop signal, with findings indicating that perceptual color similarity between Go and Stop signals increased SSRT.

While all these studies underscore the importance of perceptual discriminability in motor control, they primarily address lower-level sensory factors; our findings expand this framework by focusing on the cognitive informational content, varying the load required to extract meaning from stimuli while maintaining constant perceptual features. In our task, both the Go and Stop signals are tied to a mathematical rule requiring participants to evaluate the numerical relationship between numbers by manipulating the ND, thereby introducing a factor intervening in a phase of stimulus processing after the perceptual encoding, which we show to significantly influence motor decision. In fact, our results (Figure 2) show that when the stimulus processing engages a higher cognitive load, movement initiation is delayed, as when perceptual ambiguity is higher (Middlebrooks & Schall, 2013), but also the same factor influenced the recruitment of the Stop process. These distance-related effects have been previously observed to influence motor decisions (Acuna et al., 2002; Brunamonti et al., 2012, 2016; Krajcsi et al., 2016; Mione et al., 2020; Ramawat et al., 2022, 2023; for a review), and here we provide evidence that they also intervene in modulating inhibitory control. Specifically, participants’ performance during movement initiation showed improved accuracy and faster RTs at larger NDs, indicating that more distinct numerical contrasts reduce cognitive demand and facilitate decision making. Furthermore, the hypothesis of a spatially-oriented representation was supported by the fact that RTs were modulated by the SL of the Go signals so that when higher numbers appeared on the right side, responses were faster than when they appeared on the Left. Similarly, ND also modulated movement inhibition in Stop trials (Figure 3). Within the SST framework, which models movement inhibition as the outcome of a race between independent Go and Stop processes (Logan & Cowan, 1984), we observed that p(Respond) depends on the time the Stop process is recruited and how far the Go process is from the movement’s onset threshold. As typically described here, we observed that SSD is a factor influencing motor inhibition, but here we provide evidence that variables interact with the difficulty in engaging the Stop process as a function of the ND. It is likely that the time required to process closer numerical values delayed the initiation or progression of the Stop process, delaying the SSRT, which we observed to be longer for smaller ND, allowing the Go response to reach threshold. In addition, more participants failed to meet the model’s independence assumptions criteria at smaller NDs, indicating that violations were themselves ND-dependent and more common under higher cognitive load. Although the independence assumption is typically met in classical SST studies (Verbruggen & Logan, 2009a), previous research has shown that dependence between Go and Stop processes can emerge in selective SST variants, especially when response selection becomes more complex (De Jong et al., 1995; Bissett & Logan, 2014). Verbruggen and Logan (2015) similarly reported increased dependence when the Stop signal was harder to discriminate or interpret. By comparing mean failed Stop RTs with mean Go RTs, they found that, under higher signal ambiguity (varied-mapping conditions), failed Stop RTs were longer than Go RTs, indicating a violation of independence. In contrast, failed Stop RTs remained shorter in low-ambiguity (consistent-mapping conditions), consistent with race model predictions. Our results align with this view, showing that increased decision difficulty at smaller NDs was associated with greater dependence between processes, as reflected in a higher number of participants exhibiting longer mean Stop Error RTs than mean Go RTs.

Extending our behavioral findings, the modeling results provided deeper insights into how cognitive interpretability shapes motor decision-making during movement initiation (Figure 4 **b-c**). Using the GDDM, which decomposes the decision process by isolating key parameters such as drift rate, decision boundary, and starting point, we were able to identify the latent cognitive parameters that underlie movement initiation during Go trials. In our task design, we assumed a fixed starting point due to the simple button-release response mapping, which enabled us to control for internal factors like reward expectancy or motivation that are known to alter it (Ratcliff, 1985; Diederich & Busemeyer, 2006; Giuffrida et al., 2023). Similar to this, we assumed stable decision boundaries because the task did not include explicit speed-accuracy trade-offs or perceptual uncertainty, ruling out variations in response caution or strategy (Ratcliff et al., 2015). The best-fitting model (Model 4) among the tested models had ND-modulated Go Drifts, an SL bias, and a leak parameter that showed imperfect integration over time. This result is consistent with classical effects that indicate slower and more error-prone comparisons are produced by lower ND (Moyer & Landauer, 1967; Krajcsi et al., 2016). This suggests that the drift rate mainly reflects cognitive demand. In parallel, the spatial configuration of the response targets introduced an SL bias. Specifically, trials where the higher number appeared on the right side consistently produced higher drift rates, suggesting a spatial advantage. This SL bias, consistent with the MNL, proposes a left-to-right spatial mapping of numerical values and is consistent with prior findings on spatial-numerical associations influencing movement selection and perceptual judgments (Dehaene et al., 1993; Umiltà et al., 2009).

In our final analysis, using a GLMM analysis, we examined the influence of previously analyzed variables on successful movement inhibition. We observed that a higher Go Drift (i.e., faster evidence accumulation in the Go process) had the strongest influence among the variables analyzed, consistent with findings that higher response preparation can facilitate movement initiation but hinder motor control (Andujar et al., 2022). The second most influential variable in action stopping was the duration of the SSD. Both variables, whether considered individually or in interaction, align with the core predictions of the race model (Logan & Cowan, 1984). The GLMM also indicated that other variables, such as Stop ND, modulated these effects, particularly when the Stop signal was easy to decode. In such cases, longer SSDs amplified the impact of high Go Drifts, while higher Stop ND mitigated it, increasing the likelihood of inhibition.

These findings indicate that movement inhibition depends not only on the timing of the Stop signal but also on how effectively participants extract and process relevant information content from both Go and Stop signals. At the single-trial level, the probability of interrupting an undesired movement reflects the configuration of multiple interacting variables. Specifically, successful movement inhibition is more likely when the Go process has a lower Go Drift rate, the Stop signal appears early (short SSD), and the Stop signal is easily encoded (high Stop ND). On the contrary, inhibition is impaired when these variables assume opposite values. Overall, our results underscore the importance of cognitive variables involved in processing the Stop signal and how their interaction with core SST variables, such as Go RT and SSD (Logan & Cowan, 1984), shapes movement inhibition.

These findings are consistent with prior work emphasizing the role of intervening variables in shaping the race dynamics between Go and Stop processes (Fiori et al., 2025; Ramawat et al., 2024), and they extend the set of variables that should be considered when applying the SST in studying motor inhibition across different contexts.

## Author contributions

E.B. and S.Fe. supervised the overall project. E.B., S.Fe., and S.Fa. provided key resources and research infrastructure. E.B. and I.B.M. conceived and designed the study, carried out the investigation, and contributed to data interpretation. I.B.M. developed the methodology, performed formal analysis, and prepared the visualizations and figures. I.B.M. and V.G. conducted the modeling and simulation work. M.S. and A.P. were responsible for data acquisition, curation, and recruitment of participants for the study. E.B., I.B.M., and V.G. wrote the original draft of the manuscript. All authors reviewed, edited, and approved the final version of the manuscript.

## REFERENCES

Acuna, B. D. (2002). Frontal and Parietal Lobe Activation during Transitive Inference in Humans. Cerebral Cortex, 12(12), 1312–1321. 10.1093/cercor/12.12.1312

Andujar, M., Marc, I. B., Giuffrida, V., Ferraina, S., Brunamonti, E., & Pani, P. (2022). Response Preparation Affects Cognitive Motor Control. Human Factors, 66(4), 975–986. 10.1177/00187208221132749

Aron, A. R., & Verbruggen, F. (2008). Stop the Presses Dissociating a Selective from a Global Mechanism for Stopping. Psychological Science, 19, 1146–1153. https://www.psychtoolbox.org

Bissett, P. G., & Logan, G. D. (2011). Balancing cognitive demands: Control adjustments in the stop-signal paradigm. Journal of Experimental Psychology: Learning Memory and Cognition, 37(2), 392– 404. 10.1037/a0021800

Bissett, P. G., & Logan, G. D. (2014). Selective Stopping? Maybe Not. Joumal of Experimental Psychology: General, 143(1), 455–472. 10.1037/aOO32122

Boehler, C. N., Hopf, J. M., Stoppel, C. M., & Krebs, R. M. (2012). Motivating inhibition - reward prospect speeds up response cancellation. Cognition, 125(3), 498–503. 10.1016/j.cognition.2012.07.018

Boehler, C. N., Schevernels, H., Hopf, J. M., Stoppel, C. M., & Krebs, R. M. (2014). Reward prospect rapidly speeds up response inhibition via reactive control. Cognitive, Affective and Behavioral Neuroscience, 14(2), 593–609. 10.3758/s13415-014-0251-5

Boucher, L., Palmeri, T. J., Logan, G. D., & Schall, J. D. (2007). Inhibitory control in mind and brain: An interactive race model of countermanding saccades. Psychological Review, 114(2), 376–397. 10.1037/0033-295X.114.2.376

Brunamonti, E., Falcone, R., Genovesio, A., Costa, S., & Ferraina, S. (2012). Gaze orientation interferes with mental numerical representation. Cognitive Processing, 13, 375–379 (2012). 10.1007/s10339-012-0517-1

Brunamonti, E., Mione, V., di Bello, F., Pani, P., Genovesio, A., & Ferraina, S. (2016). Neuronal modulation in the prefrontal cortex in a transitive inference task: Evidence of neuronal correlates of mental schema management. Journal of Neuroscience, 36(4), 1223–1236.

De Jong, R., Coles, M. G., & Logan, G. D. (1995). Strategies and mechanisms in nonselective and selective inhibitory motor control. Journal of Experimental Psychology: Human Perception and Performance, 21(3), 498–511. 10.1037/0096-1523.21.3.498

Dehaene, S. (1997). The number sense: how the mind creates mathematics. Oxford University Press, Inc., 1997.

Dehaene, S., Bossini, S., & Giraux, P. (1993). The Mental Representation of Parity and Number Magnitude. Journal of Experimental Psychology:General, 122(3).

Diederich, A., & Busemeyer, J. R. (2006). Modeling the effects of payoff on response bias in a perceptual discrimination task: Bound-change, drift-rate-change, or two-stage-processing hypothesis. Perception & Psychophysics, 68(2), 194–207. 10.3758/BF03193669

Dix, A., & Li, S. C. (2020). Incentive motivation improves numerosity discrimination: Insights from pupillometry combined with drift-diffusion modelling. Scientific Reports, 10(1). 10.1038/s41598-020-59415-3

Fischer, M. H., & Shaki, S. (2014). Spatial associations in numerical cognition-From single digits to arithmetic. The Quarterly Journal of Experimental Psychology, 67(8), 1461–1483. 10.1080/17470218.2014.927515

Fiori, L., Ramawat, S., Marc, I.B., Giuffrida, V., Ranavolo, A., Draicchio, F., Pani, P., Ferraina, S., Brunamonti, E. (2025). Balancing postural control and motor inhibition during gait initiation. iScience 28(3), 2589–0042; 10.1016/j.isci.2025.111970

Giamundo, M., Giarocco, F., Brunamonti, E., Fabbrini, F., Pani, P., & Ferraina, S. (2021). Neuronal Activity in the Premotor Cortex of Monkeys Reflects Both Cue Salience and Motivation for Action Generation and Inhibition. Journal of Neuroscience, 41(36) 7591–7606; 10.1523/JNEUROSCI.0641-20.2021

Giuffrida, V., Marc, I. B., Ramawat, S., Fontana, R., Fiori, L., Bardella, G., Fagioli, S., Ferraina, S., Brunamonti, E., & Pani, P. (2023). Reward prospect affects strategic adjustments in stop signal task. Frontiers in Psychology, 14. 10.3389/fpsyg.2023.1125066

Haque, M.T., Segreti, M., Giuffrida, V.S., Ferraina, S., Brunamonti, E., & Pani, P. (2024). Attentional spatial cueing of the stop-signal affects the ability to suppress behavioural responses. Experimental Brain Research, 242, 1429–1438. 10.1007/s00221-024-06825-8

Hilt PM, Cardellicchio P (2020) Attentional bias on motor control: is motor inhibition influenced by attentional reorienting? Psychological Research, 84(2), 276–284. 10.1007/s00426-018-0998-3

Holloway, I. D., & Ansari, D. (2008). Mapping numerical magnitudes onto symbols: The numerical distance effect and individual differences in children’s mathematics achievement. Journal of Experimental Child Psychology, 103(1), 17–29. 10.1016/j.jecp.2008.04.001

Hubbard, E. M., Diester, I., Cantlon, J. F., Ansari, D., Van Opstal, F., & Troiani, V. (2008). The evolution of numerical cognition: From number neurons to linguistic quantifiers. Journal of Neuroscience, 28(46), 11819–11824. 10.1523/JNEUROSCI.3808-08.2008

Hubbard, E. M., Piazza, M., Pinel, P., & Dehaene, S. (2005). Interactions between number and space in parietal cortex. In Nature Reviews Neuroscience (Vol. 6, Issue 6, pp. 435–448). 10.1038/nrn1684

Izard, V., & Dehaene, S. (2008). Calibrating the mental number line. Cognition, 106(3), 1221–1247. 10.1016/j.cognition.2007.06.004

Jensen, G., Alkan, Y., Muñoz, F., Ferrera, V. P., & Terrace, H. S. (2017). Transitive inference in humans (Homo sapiens) and rhesus macaques (Macaca mulatta) after massed training of the last two list items. Journal of Comparative Psychology, 131(3), 231–245. 10.1037/com0000065

Krajcsi, A., Lengyel, G., & Kojouharova, P. (2016). The source of the symbolic numerical distance and size effects. Frontiers in Psychology, 7(NOV). 10.3389/fpsyg.2016.01795

Leotti, L. A., & Wager, T. D. (2010). Motivational Influences on Response Inhibition Measures. Journal of Experimental Psychology: Human Perception and Performance, 36(2), 430–447. 10.1037/a0016802

Logan, G. D. (1981). Attention and performance IX. In Attention, automaticity, and the ability to stop a speeded choice response (pp. 205–222).

Logan, G. D., & Cowan, W. B. (1984). On the Ability to Inhibit Thought and Action: A Theory of an Act of Control. In Psychological Review (Vol. 91, Issue 3).

Logan, G. D., & Irwin, D. E. (2000). Don’t look! Don’t touch! Inhibitory control of eye and hand movements. In Psychonomic Bulletin & Review (Vol. 7, Issue I).

Longo, M. R., & Lourenco, S. F. (2007). Spatial attention and the mental number line: Evidence for characteristic biases and compression. Neuropsychologia, 45(7), 1400–1407. 10.1016/j.neuropsychologia.2006.11.002

Marc, I. B., Giuffrida, V., Ramawat, S., Fiori, L., Fontana, R., Bardella, G., Fagioli, S., Ferraina, S., Pani, P., & Brunamonti, E. (2023). Restart errors reaction time of a two-step inhibition process account for the violation of the race model’s independence in multi-effector selective stop signal task. Frontiers in Human Neuroscience, 17. 10.3389/fnhum.2023.1106298

Merritt, D. J., & Terrace, H. S. (2011). Mechanisms of Inferential Order Judgments in Humans (Homo sapiens) and Rhesus Monkeys (Macaca mulatta). Journal of Comparative Psychology, 125(2), 227–238. 10.1037/a0021572

Middlebrooks, P. G., & Schall, J. D. (2013). Response inhibition during perceptual decision making in humans and macaques. Attention, Perception, and Psychophysics, 76(2), 353–366. 10.3758/s13414-013-0599-6

Middlebrooks, P. G., Zandbelt, B. B., Logan, G. D., Palmeri, T. J., & Schall, J. D. (2019). Countermanding Perceptual Decision-Making. IScience, 23(1). 10.1016/j.isci.2019.100777

Middlebrooks, P. G., Zandbelt, B. B., Logan, G. D., Palmeri, T. J., Schall, J. D., Feurtado, E. M., Maddox, M., Motorny, S., Parker, J., & Schall, M. (2017). Unification of Countermanding and Perceptual Decision-making. BioRxiv. 10.1101/158170

Mione, V., Brunamonti, E., Pani, P., Genovesio, A., & Ferraina, S. (2020). Dorsal Premotor Cortex Neurons Signal the Level of Choice Difficulty during Logical Decisions. Cell Reports, 32(4). 10.1016/j.celrep.2020.107961

Montanari, R., Giamundo, M., Brunamonti, E., Ferraina, S., & Pani, P. (2017). Visual salience of the stop-signal affects movement suppression process. Experimental Brain Research, 235(7), 2203–2214. 10.1007/s00221-017-4961-0

Morein-Zamir, S., & Kingstone, A. (2006). Fixation offset and stop signal intensity effects on saccadic countermanding: A crossmodal investigation. Experimental Brain Research, 175(3), 453–462. 10.1007/s00221-006-0564-x

Moyer, R. S., & Landauer, T. K. (1967). Time required for Judgements of Numerical Inequality. Nature, 215, 1519–1520.

Oldfield, R. C. (1971). THE ASSESSMENT AND ANALYSIS OF HANDEDNESS: THE EDINBURGH INVENTORY. In Neuropsychologia (Vol. 9). Pergamon Press.

Osman, A., Kornblum, S., & Meyer, D. E. (1986). The Point of No Return in Choice Reaction Time: Controlled and Ballistic Stages of Response Preparation. In Journal of Experimental Psychology: Human Perception and Performance (Vol. 12, Issue 3).

Pani, P., Giarrocco, F., Giamundo, M., Montanari, R., Brunamonti, E., & Ferraina, S. (2018). Visual salience of the stop signal affects the neuronal dynamics of controlled inhibition. Scientific Reports, 8(1). 10.1038/s41598-018-32669-8

Park, J., & Starns, J. J. (2015). The approximate number system acuity redefined: A diffusion model approach. Frontiers in Psychology, 6(DEC). 10.3389/fpsyg.2015.01955

Ramawat, S., Marc, I. B., Ceccarelli, F., Ferrucci, L., Bardella, G., Ferraina, S., Pani, P., & Brunamonti, E. (2023). The transitive inference task to study the neuronal correlates of memory-driven decision making: A monkey neurophysiology perspective. In Neuroscience and Biobehavioral Reviews (Vol. 152). Elsevier Ltd. 10.1016/j.neubiorev.2023.105258

Ramawat, S., Marc, I. B., di Bello, F., Bardella, G., Ferraina, S., Pani, P., & Brunamonti, E. (2024). Force monitoring reveals single trial dynamics of motor control in a stop signal task. Physiological Reports, 12(22); 10.14814/phy2.70127

Ramawat, S., Mione, V., Di Bello, F., Bardella, G., Genovesio, A., Pani, P., Ferraina, S., & Brunamonti, E. (2022). Different Contribution of the Monkey Prefrontal and Premotor Dorsal Cortex in Decision Making During a Transitive Inference Task. Neuroscience, 485, 147–162. 10.1016/j.neuroscience.2022.01.013

Ratcliff, R., & Mckoon, G. (2008). The Diffusion Decision Model: Theory and Data for Two-Choice Decision Tasks. Neural Computation, 20, 873–922.

Ratcliff, R., Thompson, C. A., & McKoon, G. (2015). Modeling individual differences in response time and accuracy in numeracy. Cognition, 137, 115–136. 10.1016/j.cognition.2014.12.004

Shinn, M., Lam, N.H., & John D Murray, J.D. (2020). A flexible framework for simulating and fitting generalized drift-diffusion models. eLife 9:e56938. 10.7554/eLife.56938

Umiltà, C., Priftis, K., & Zorzi, M. (2009). The spatial representation of numbers: Evidence from neglect and pseudoneglect. Experimental Brain Research, 192(3), 561–569. 10.1007/s00221-008-1623-2

Van Der Schoot, M., Licht, R., Horsley, T. M., & Sergeant, J. A. (2005). Effects of stop signal modality, stop signal intensity and tracking method on inhibitory performance as determined by use of the stop signal paradigm. Scandinavian Journal of Psychology, 46 (4), 331–341. 10.1111/j.1467-9450.2005.00463.x

Verbruggen, F., Aron, A. R., Band, G. P. H., Beste, C., Bissett, P. G., Brockett, A. T., Brown, J. W., Chamberlain, S. R., Chambers, C. D., Colonius, H., Colzato, L. S., Corneil, B. D., Coxon, J. P., Dupuis, A., Eagle, D. M., Garavan, H., Greenhouse, I., Heathcote, A., Huster, R. J., … Boehler, C. N. (2019). A consensus guide to capturing the ability to inhibit actions and impulsive behaviors in the stop-signal task. ELife, 8. 10.7554/eLife.46323

Verbruggen, F., & Logan, G. D. (2009a). Models of response inhibition in the stop-signal and stop-change paradigms. Neuroscience & Biobehavioral Reviews, 33(5), 647–661. 10.1016/j.neubiorev.2008.08.014.

Verbruggen, F., & Logan, G. D. (2015). Evidence for capacity sharing when stopping. Cognition, 145, 81–95. 10.1016/j.cognition.2015.05.014

Verbruggen, F., & McLaren, R. (2018). Effects of reward and punishment on the interaction between going and stopping in a selective stop-change task. Psychological Research, 82(2), 353–370. 10.1007/s00426-016-0827-5

Verbruggen, F., Stevens, T., Chambers, C.D. (2014) Proactive and reactive stopping when distracted: an attentional account. Journal of Experimental Psychology: Human Perception and Performance. 40(4):1295–300. 10.1037/a0036542.

